# The strand-selective regulation of miR-155 in response to lipopolysaccharide by CELF2, FUBP1 and KSRP proteins

**DOI:** 10.1101/2025.03.05.641704

**Authors:** Jeff S. J. Yoon, Abdulwadood Baksh, Thomas C. Chamberlain, Alice L-F Mui

## Abstract

The microRNA-155 exists in two forms, miR-155-5p and miR-155-3p, produced from either strand of the double-stranded precursor of miR-155. The more abundant and better-studied miR-155-5p has been implicated in many biological processes with dysregulated expression occurring in human diseases. miR-155-5p plays an essential role in supporting inflammatory responses in macrophages. Activating macrophages with lipopolysaccharide elevates miR-155-5p, while the anti-inflammatory cytokine interleukin-10 reduces miR-155-5p levels. Recently, miR-155-3p has also been suggested to participate in macrophage function, although its specific role is unclear. We found that LPS stimulation of macrophages results first in the elevation of miR-155-3p levels, followed by an increase in miR-155-5p levels. In this paper, we explore the mechanisms involved in the maturation of the pre-miR-155 to either miR-155-5p or miR-155-3p. We describe the contribution of three RNA-binding proteins, CELF2, FUBP1 and KSRP, to pre-miR-155 processing. Our data suggests CELF2 regulates miR-155-5p/miR-155-3p strand selection. FUBP1 may support the expression of miR-155-3p for specific subcellular functions, while KSRP appears to inhibit both miR-155-5p and miR-155-3p maturation without altering the relative expression of each strand.

## Introduction

MicroRNAs (miRNAs) are short regulatory non-coding RNAs that post-transcriptionally regulate the expression of genes through sequence-specific targeting of mRNA^1^. MiR-155 is among the most extensively studied miRNAs due to its involvement in various biological and pathological processes, including immunity, inflammation and cancer^2–8^. The MIR155HG locus encodes miR-155, and the primary transcript of miR-155 (**pri-miR-155**) undergoes Drosha-mediated cleavage to the precursor of miR-155 (**pre-miR-155**)^9^. Pre-miR-155 can undergo processing into two alternate forms, the dominant strand miR-155-5p and the non-dominant strand miR-155-3p^10^, depending on which strand from the pre-miR-155 is selected. The selected strand is loaded onto the RNA-induced silencing complex (RISC) and becomes the guide strand, while the other strand undergoes degradation^11^. The 5p and 3p strands have different seed sequences and thus target different mRNAs^12^. Although miR-155-5p is the most abundant and studied strand, miR-155-3p was recently shown to have biological functions^13–15^. Furthermore, the dysregulated expression of miR-155-3p alone has been observed in certain cancers and diseases ^16–21^.

Activating macrophage cells is essential in the human host’s defence against pathogens^22,23^. We^24^ and other^3,4,25^ showed that activation of macrophage cells with lipopolysaccharide (LPS), a cell wall component of gram-negative bacteria, induced expression of miR-155-5p. miR-155-5p was the only strand studied and required for macrophage production of inflammatory cytokines such as tumour necrosis factor-alpha (TNFα). However, persistent macrophage activation can become harmful, leading to inflammatory diseases, cardiovascular diseases and cancers^26–28^. Consequently, activated macrophages also release anti-inflammatory cytokines, namely interleukin-10 (IL10), to attenuate the pro-inflammatory effects of macrophages activated by LPS^29^. IL10 is a pleiotropic cytokine characterized initially as a cytokine synthesis inhibitory factor produced by murine Th T-cells to prevent cytokine production by murine Th T-cells^29^. However, deactivating the activated macrophages is its major *in vivo* function^30^. IL10 receptor (IL10R) signalling uses both the Signal Transducer and Activator of Transcription 3 (STAT3)^31–33^ and Src Homology 2 domain-containing Inositol-5 Phosphatase 1 (SHIP1)-dependent^24,31,34^ pathways to deactivate macrophages^24,32,34–36^. We previously reported that SHIP1 could form a complex with STAT3 (SHIP1:STAT3) in response to IL10R signalling or through the exposure of cells to small molecules that bind to induce a conformational change in SHIP1^31^. The formation of the SHIP1:STAT3 complex is sufficient to inhibit macrophage production of TNFα. We also showed that IL10 inhibition of miR-155-5p required SHIP1 and STAT3^24^.

We examined how IL10 interferes with the LPS-induced expression of miR-155-5p. We found that IL10 inhibits the maturation of pre-miR-155 into mature miR-155-5p^24^. We then used mass spectrometry to identify proteins that bind to pre-miR-155 to characterize the mechanism by which IL10R signalling inhibits pre-miR-155 maturation^37^. CUGBP Elav-Like Family member 2 (CELF2) protein is an RNA-binding protein known for its post-transcriptional regulation of mRNAs through alternative splicing of mRNAs^38,39^ or polyadenylation^40^. CELF2 and its family member CELF1 also participate in microRNA regulation^41^. Treiber *et al*. showed that CELF2 and CELF1 proteins bind to the stem of pre-miR-140 and inhibit its maturation to miR-140^41^. In our previous study, we found that CELF2 protein associates with pre-miR-155 in an IL10-dependent manner, and deletion of CELF2 enhanced LPS-induced expression of miR-155-5p, suggesting CELF2’s role in inhibiting miR-155-5p expression^37^.

Recently, Simmonds *et al.* showed that LPS stimulation also leads to the early expression of miR-155-3p, followed by miR-155-5p at later time points^42^. Thus, in the current study, we examined whether CELF2 and two other proteins we identified to bind pre-miR-155, Far Upstream Element Binding Protein 1 (**FUBP1**) and KH-type Splicing Regulatory Protein (**KSRP**)^37,43^, might be involved in regulating miR-155-3p and miR-155-5p levels, especially the switch from expressing the miR-155-3p to the miR-155-5p strand. FUBP1 belongs to a family of RNA-binding proteins (FUBP family), including KSRP^43^. Previously, studies implicated KSRP in controlling miR-155-5p expression in macrophages^25,44^ and dendritic cells^45^. Ruggiero *et al.* reported that pre-miR-155 co-immunoprecipitated with KSRP and depletion of KSRP impaired the expression of mature miR-155-5p while simultaneously causing the accumulation of pri-miR-155 and pre-miR-155^25^. KSRP, also known as Far Upstream Element Binding Protein 2 (FUBP2), binds to the terminal loop of pre-miR-155 via four distinct hnRNPK-Homology (KH) domains and promotes its maturation^25,46,47^.

We now describe the roles of CELF2, FUBP1 and KSRP in controlling the expression of miR-155-5p and 3p. We used CRISPR-Cas9 mediated silencing to construct cells deficient in CELF2, FUBP1, or KSRP. Our data suggest that CELF2 is involved in the timing of miR-155-3p expression and affects the overall strand ratio of miR-155-3p and miR-155-5p. FUBP1 specifically inhibits miR-155-3p, while KSRP inhibits the expression of both strands. Finally, we show that FUBP1 and KSRP proteins interact with pre-miR-155 via its KH3 domain. These data provide insights into dysregulated miR-155-5p and miR-155-3p observed in certain diseases.

## Results

### LPS induces the association of FUBP1 to pre-miR-155

We and others have shown that IL10 inhibits LPS-induced miR-155-5p expression in macrophage cells^24,37^. We further showed that IL10 inhibits the maturation of pre-miR-155 to miR-155-5p^24^ and that the RNA-binding protein CELF2 contributes to the process^37^. However, our mass spectrometry-based examination of pre-miR-155-associated proteins also identified FUBP1 and KSRP as other proteins that might interact with pre-miR-155 in a stimulus (LPS ± IL10) dependent manner^37^.

We analyzed pre-miR-155 pull-down samples for CELF2, FUBP1 and KSRP protein to follow up on the mass spectrometry identified protein candidates. As described in our previous study^37^, the amount of CELF2 detected in the pre-miR-155 pull-down samples is the greatest in the LPS + IL10 samples (**Fig. 1**). LPS stimulation of the cells also increases the amount of FUBP1 protein in the pre-miR-155 pull-down samples; the addition of IL10 increases this further. KSRP levels in the pre-miR-155 pull-down samples appear to rise with LPS and further with IL10 addition. Still, these changes are not statistically significant (**Fig. 1**). The levels of all three proteins in the total cell lysate remained constant in all stimulation conditions, suggesting that the association of these proteins with pre-miR-155 might represent post-transcriptional modifications of those proteins, which alters their ability to bind pre-miR-155.

**Fig. 1.**
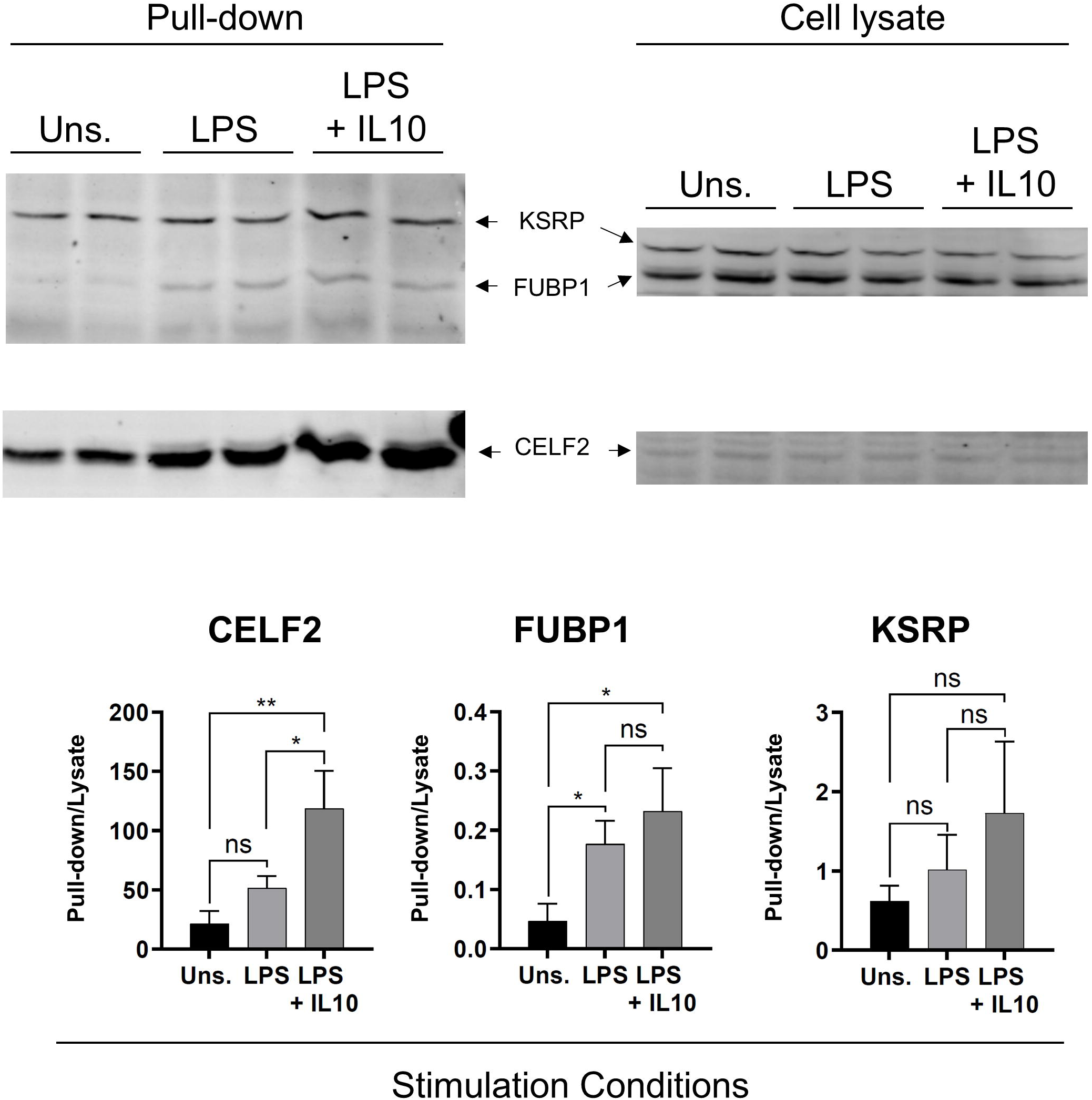
Increased interaction of FUBP1 protein with pre-miR-155 in response to LPS stimulation. RAW264.7 cells were transfected with biotinylated pre-miR-155 oligonucleotide and stimulated with LPS ± IL10 for 2 hours before collecting the pull-down samples. Expression levels of CELF2, FUBP1 and KSRP protein interacting with pre-miR-155 oligonucleotide were determined by immunoblotting. The graph shows the CELF2, FUBP1 and KSRP band intensities in the pull-down sample normalized to the same protein in total cell lysate. One-way ANOVA with Tukey’s correction calculated the significance of the difference between the stimulations. ** *p* < 0.01, * *p* < 0.05, ns = not significant. The data are representative of 5 experiments.

### CELF2, FUBP1 and KSRP deficiency differentially affects miR-155-5p and miR-155-3p expression

We^24^ and others have shown that IL10 reduces miR-155-5p expression in LPS-stimulated macrophages^24,25,37^. To investigate the role of CELF2, FUBP1 and KSRP in LPS and IL10 action in these pathways, we used CRISPR-Cas9-mediated targeting to knock down the expression of each of the three proteins in RAW264.7 cells. We transduced CRISPR-Cas9 expressing RAW264.7 cells with lentiviruses harbouring a Blasticidin resistance gene and a guide RNA (gRNA) sequence targeting either FUBP1 or KSRP, as we described previously for CELF2 KD cells^37^. Cells were selected with 10 µg/mL Blasticidin, and the bulk drug-resistant population was used for experiments. We call CRISPR-Cas9/gRNA transduced cells “KD” for knockdown rather than KO (for knockout) because we use a bulk, uncloned population, which may contain cells with different degrees of target gene deletion. We use a bulk population to reduce the impact of insertional mutagenesis effects, which the use of cell clones would magnify. As illustrated in **Fig. 2**, FUBP1 KD and KSRP KD cells express significantly less FUBP1 or KSRP protein than the control RAW264.7 cell.

**Fig. 2.**
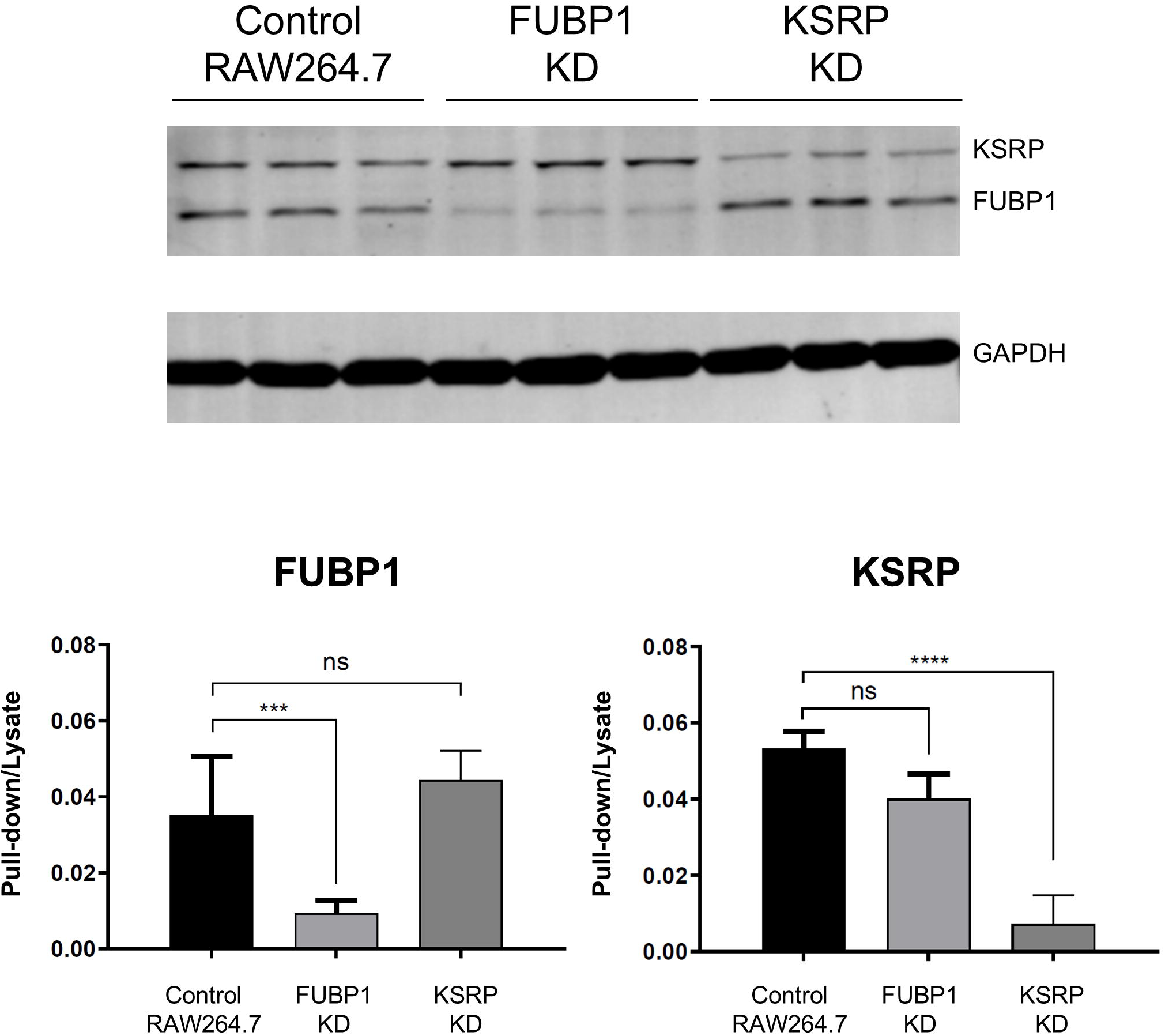
Generation of FUBP1 and KSRP knockdown cells by CRISRP-Cas9 mediated gene silencing. RAW264.7/Cas9 cells transduced with FUBP1 sgRNA or KSRP sgRNA were treated with 2 µg/mL of doxycycline for 48 hours to induce knockdown of FUBP1/KSRP proteins. FUBP1 and KSRP protein expression was determined by immunoblotting. Data plotted represents FUBP1 and KSRP protein band intensity normalized to GAPDH. One-Way ANOVA with Tukey’s correction determined the comparison indicated with braces. **** *p* < 0.0001, *** *p* < 0.001, ns = not significant. The data are representative of 3-5 experiments.

After confirming the reduction of FUBP1 and KSRP proteins in the KD cells, we stimulated the cells with LPS ± IL10 for 0, 2 and 4 hours, isolated RNA, and quantified the levels of miR-155-5p and miR-155-3p by qPCR. **Fig. 3A** shows every cell’s data normalized to the LPS-stimulated sample (2 hours) of the control RAW264.7 cells, allowing comparison among all the cells. **Fig 3B** graphs the same data in a time-course format for each cell type. **Fig 3B** shows that in RAW264.7 cells, the LPS-stimulated miR-155-3p levels are detectable at 2 hours and plateaus by 4 hours. In contrast, the LPS-stimulated miR-155-5p level significantly increases only by 4 hours and continues to rise past 4 hours (data not shown). As we described earlier^37^, IL10 inhibited the expression of miR-155-5p at 4 hours but not at 2 hours. IL10 inhibited the expression of miR-155-3p at 4 hours (**Fig 3A** and **Fig 3B**), but the inhibition at 2 hours was statistically significant when the comparison was within just the RAW264.7 cells (**Fig 3B**).

**Fig. 3.**
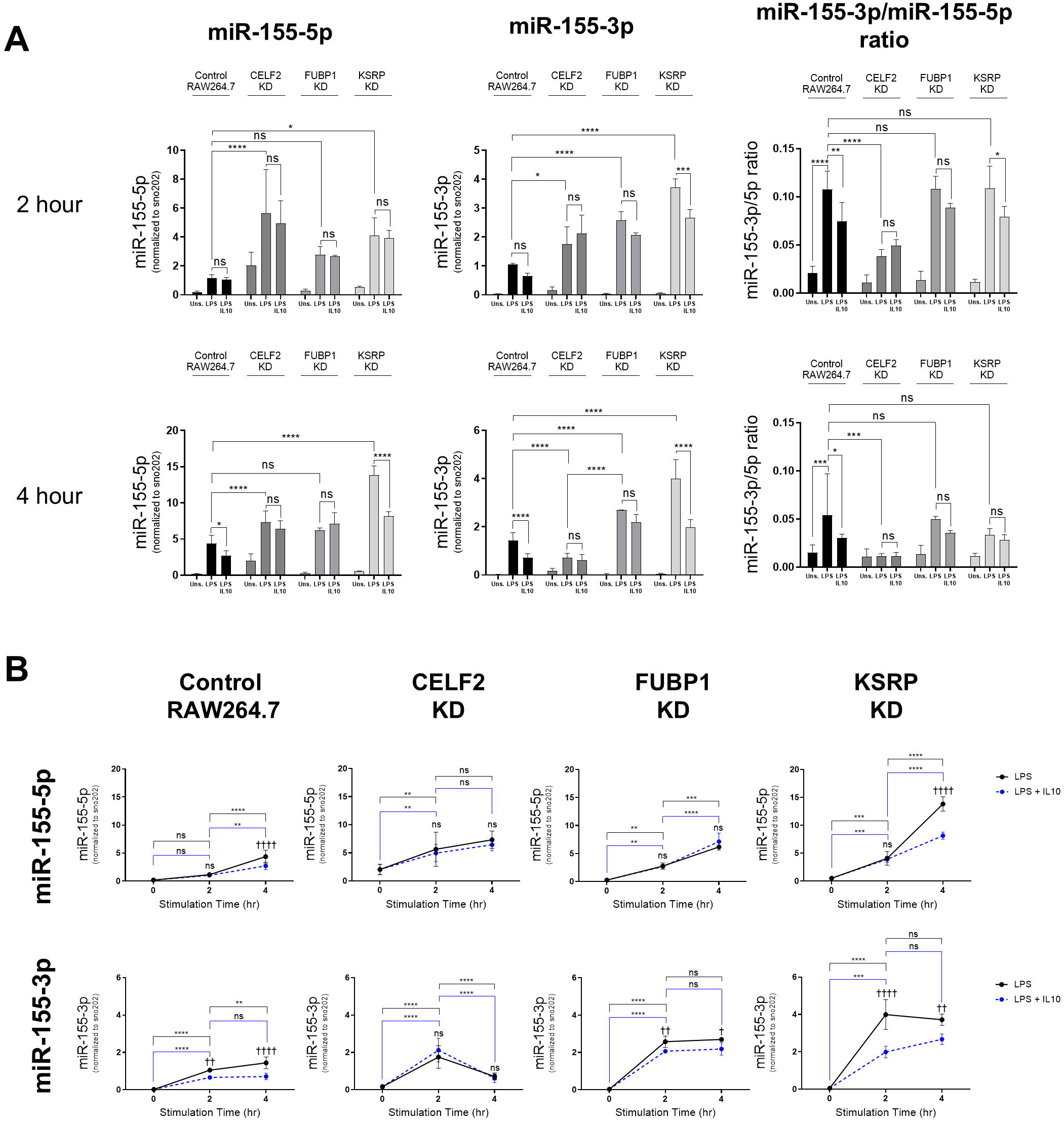
CELF2, FUBP1 and KSRP deficiency alters the expression of miR-155-5p and miR-155-3p in response to LPS ± IL10 stimulation. FUBP1 KD, CELF2 KD, KSRP KD or the control RAW264.7 cells were stimulated with 1 ng/mL LPS ± 1 ng/mL of IL10 for (**A**) 2 and 4 hours before total RNA extraction. The expression levels of miR-155-5p and miR-155-3p were determined by qPCR and normalized to snoRNA202 levels. The data plotted represents the expression levels of miR-155-5p and miR-155-3p levels normalized to the 2-hour LPS-stimulated control RAW264.7 cells. (**B**) The data from panel A were replotted as time courses for each cell type. miR-155-5p and miR-155-3p levels were normalized to the unstimulated sample of control RAW264.7 cells. Two-way ANOVA with Tukey’s correction determined the comparisons indicated with braces between different timepoints or cell type as * and the significance between stimulations as †. **** p < 0.0001, *** p < 0.001, ** p < 0.01, * p < 0.05, †††† p < 0.0001, ††† p < 0.001, †† p < 0.01, † p < 0.05, ns = not significant. The data are representative of 4 experiments.

Plotting the ratio of miR-155-3p to miR-155-5p helps illustrate the change in the selection of miR-155-3p and miR-155-5p strands for expression. In RAW264.7 cells, LPS treatment increases the ratio of miR-155-3p to miR-155-5p (**Fig. 3A**, rightmost graph), suggesting a strand selection of miR-155-3p strand over miR-155-5p. The miR-155-3p to miR-155-5p ratio is unchanged by IL10, suggesting IL10 controls miR-155 expression by inhibiting the general miR-155 maturation without altering the miR-155 strand selection. The LPS effect on the miR-155-3p to miR-155-5p ratio disappears by 4 hours.

Knockdown of CELF2 protein enhanced miR-155-5p expression at 2 and 4 hours (**Fig. 3A**) compared to control RAW264.7 cells. miR-155-3p expression is slightly enhanced in CELF2 KD compared to RAW264.7 cells at 2 hours but reduced at 4 hours of LPS stimulation (**Fig. 3A**). Of note, CELF2 KD cells show a dramatic acceleration in the kinetics of miR-155-5p and miR-155-3p expression. Compared to the control and other cells, miR-155-5p expression in CELF2 KD cells appears to have already reached a plateau by 4 hours (**Fig 3B**). miR-155-3p expression peaked earlier at 2 hrs and decreased to basal expression level by 4 hours. Interestingly, the miR-155-3p/miR155-5p ratio in CEL2 KD at 2 and 4 hours (**Fig. 3A**, right panel) significantly decreased compared to the control RAW264.7 cells. IL10 failed to inhibit the expression of either miR-155-5p or miR-155-3p in the CELF2 KD cells. The LPS-stimulated miR-155-3p/miR-155-5p ratio stays the same regardless of the addition of IL10.

In FUBP1 KD cells, LPS-stimulated miR-155-5p expression is unchanged compared to its action in control RAW264.7 cells (**Fig. 3A**). In contrast, the level of miR-155-3p is enhanced, suggesting a selective role of FUBP1 in regulating miR-155-3p expression, but the overall ratio of miR-155-3p/miR-155-5p is unchanged. Like that observed in CELF2 KD cells, IL10 could not inhibit either miR-155-5p or miR-155-3p in FUBP1 KD cells. In the KSRP KD cells, miR-155-5p and miR-155-3p expression is significantly enhanced without a change in the relative expression. This elevation occurs without significant changes in pri-miR-155 or pre-miR-155 levels (**Supplementary Fig. 1**). Unlike that observed in FUBP1 and CELF2 KD cells, IL10 could inhibit miR-155-5p and miR-155-3p in cells lacking KSRP.

### Expression of miR-155 target genes in CELF2, FUBP1 and KSRP KD cells

To assess the biological relevance of the changes in miR-155-5p and miR-155-3p changes., we examined the levels of Arg2, (a validated miR-155-5p target)^48,49^; and Apbb2 ( a predicted target of miR-155-3p) ^50,51^ and Myd88, (a target of both miR-155 strands) ^52,53^. The expression of the miR-155 targets should be decreased in cells where miR-155 levels are increased compared to RAW264.7 cells and conversely increased in cells where miR-155 levels are decreased. As expected, Arg2 levels are lower in the CELF2 and KSRP KD cells (**Fig 4**), where miR-155-5p is higher (**Fig 3A**, 4-hour data). Apbb2 levels are higher (**Fig 4**) in CELF2 KD cells, where miR-155-3p levels are lower (**Fig 3A**, 4-hour data) than in RAW264.7 cells. As expected, myd88 levels are lower in all three KD cell types (**Fig 4**) since either miR-155-5p or miR-155-3p strands are elevated in all three protein KD cells(**Fig 3A**).

**Fig. 4.**
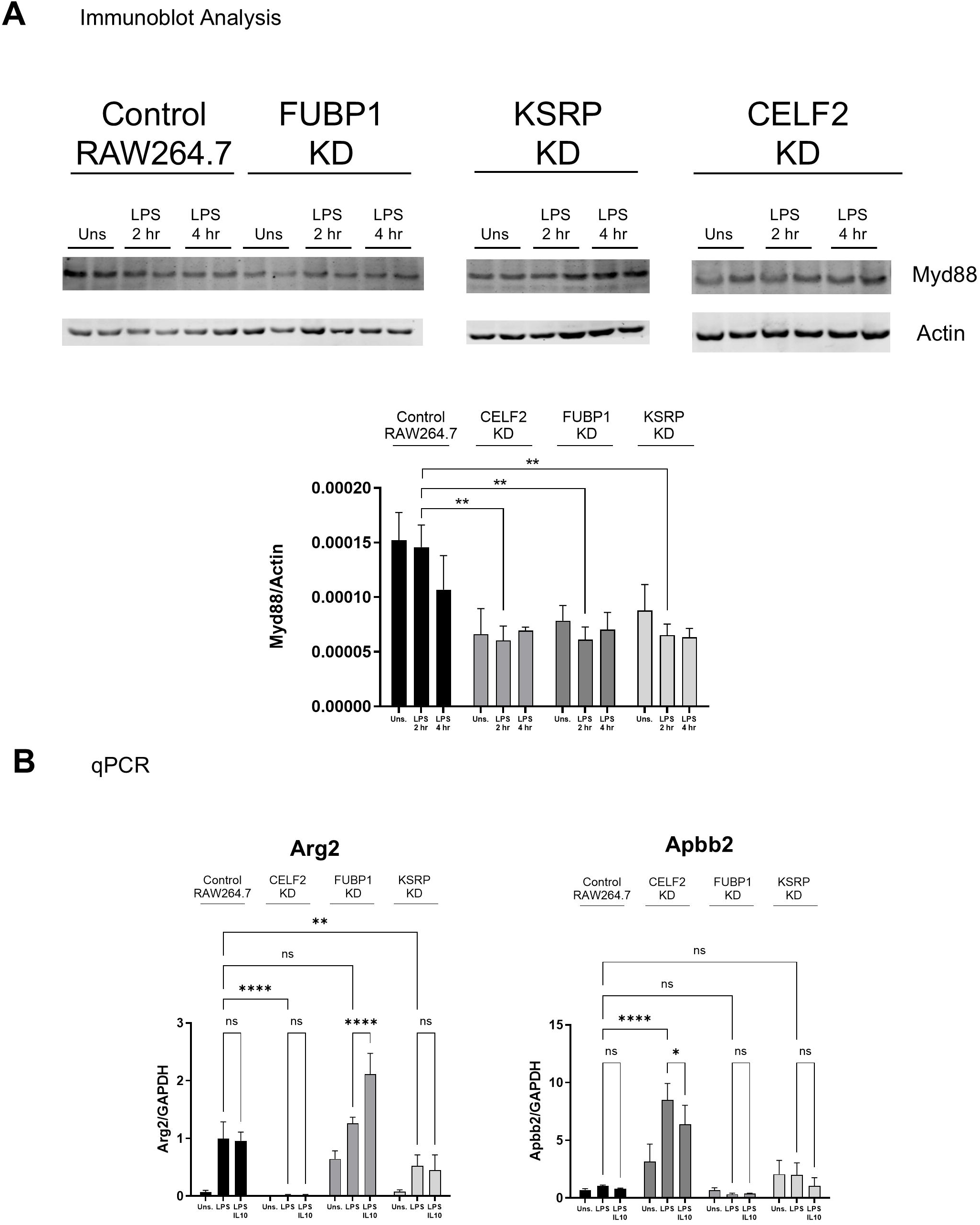
Expression of miR-155 target genes in CELF2 KD, FUBP1 KD, KSRP KD, or the control RAW264.7 cells. Cells were stimulated with 1 ng/mL of LPS 4 hours before total cell extraction. The expression levels of Arg2, Apbb2 and Myd88 were determined by qPCR and normalized to GAPDH levels. Two-way ANOVA with Tukey’s correction determined the comparisons indicated with braces. **** *p* < 0.0001, *** *p* < 0.001, ** *p* < 0.01, * *p* < 0.05, ns = not significant. The data are representative of 3 experiments.

### IL10 regulation of inflammatory cytokine production in CELF2, FUBP1, and KSRP KD cells

Previously, we and others reported that miR-155 (miR-155-5p was assessed) deficiency increases LPS-induced inflammatory cytokine production^25,37^. Therefore, we examined IL10’s action on TNFα production in FUBP1 and KSRP KD cells. Cells were stimulated with LPS ± 0.5 ng/mL or 10 ng/mL IL10 for 1 hour, and TNFα levels in the culture supernatants were quantified by ELISA. **Fig. 5** shows that in control RAW264.7 cells, IL10 inhibits LPS-induced TNFα expression in a dose-dependent manner. However, in FUBP1 KD cells, IL10 could not decrease LPS-induced TNFα levels at all the concentrations of IL10 tested. This impairment of IL10 action suggests FUBP1 is essential for IL10 inhibition of LPS-induced TNFα. We previously showed that CELF2 deficiency partially impaired IL10-dependent TNFα inhibition^37^. In contrast, the absence of KSRP protein resulted in higher LPS-stimulated TNFα production than in RAW264.7 cells, but IL10 could inhibit TNFα levels.

**Fig. 5.**
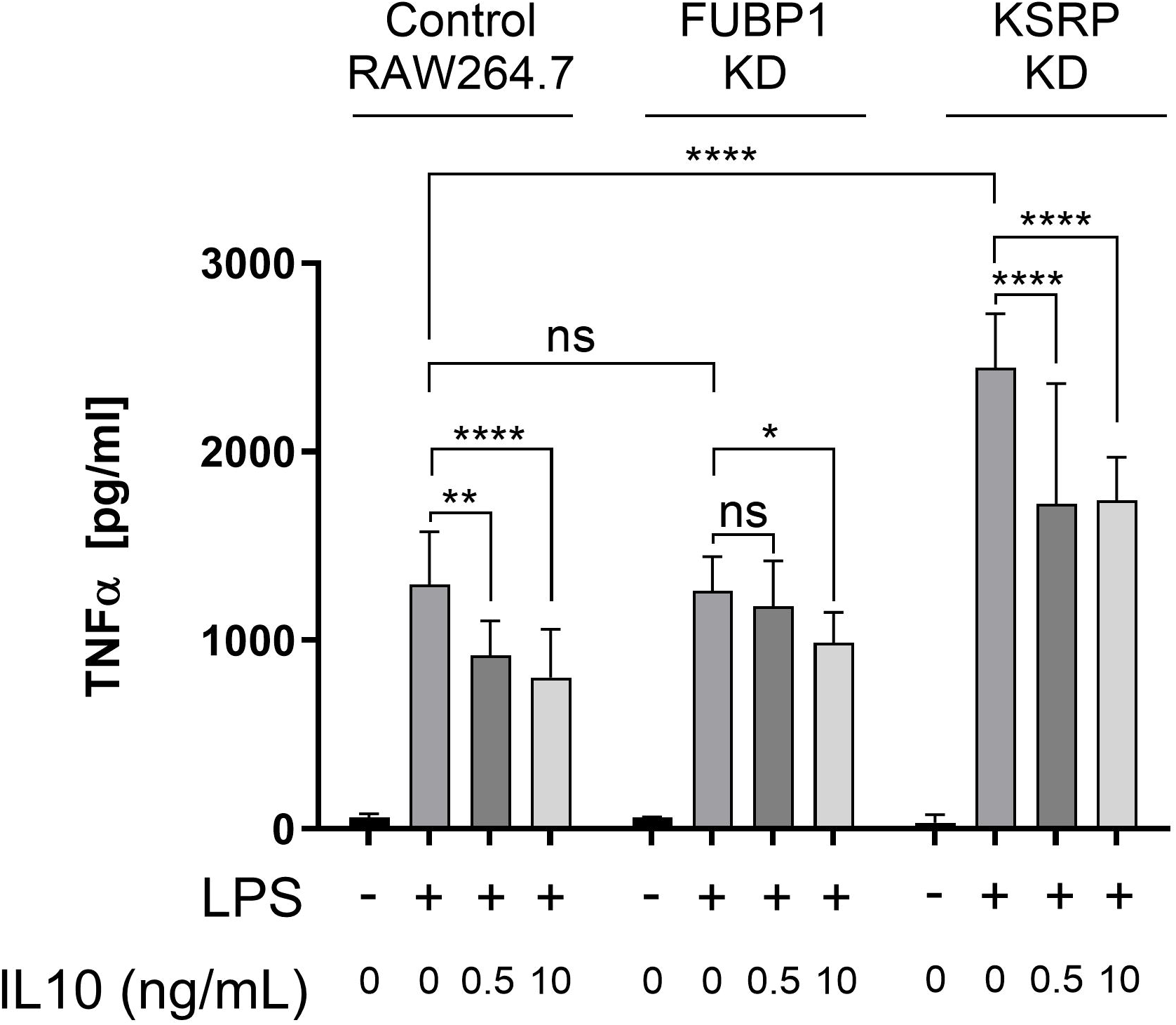
FUBP1 and KSRP deficiency alters the expression of TNFα in response to LPS ± IL10. FUBP1 and KSRP KD cells were stimulated with 1 ng/mL LPS ± indicated concentration of IL10 for 1 hour before collecting the cell culture supernatant. The level of TNFα in the supernatants was determined by ELISA. Two-Way ANOVA with Tukey’s correction determined the comparisons indicated with braces. **** *p* < 0.0001, *** *p* < 0.001, ** *p* < 0.01, * *p* < 0.05, ns = not significant. The data are representative of 3-5 experiments.

### SHIP1 and STAT3 dependence of miR-155-5p and miR-155-3p expression

We previously showed that IL10 receptor signalling requires SHIP1 and STAT3 to inhibit miR-155-5p production in LPS-stimulated macrophages^24,34^. However, we had not examined the expression of the alternate miR-155 strand, miR-155-3p. Thus, we extracted peritoneal macrophages (perimacs) from SHIP1 or STAT3 wild type (WT) or knockout (KO) mice, rested the cells for 2 hours, stimulated the cells with LPS ± IL10 for 4 hours, isolated RNA, and quantified the levels of miR-155-5p and miR-155-3p.

**Fig. 6A** shows the miR-155-5p and miR-155-3p expression in perimacs from STAT3 WT/KO (both C57Bl6 background) and SHIP1 WT/KO (both Balb/C background) mice 4 hours after LPS ± IL10 stimulation. The qPCR data were normalized to the LPS-stimulated WT sample to compare the expression in WT and KO cells. In STAT3 WT and KO perimacs, the basal and LPS-stimulated levels of miR-155-3p are similar in the corresponding sample of each cell type (**Fig. 6A**). However, LPS-stimulated levels of miR-155-5p are reduced in the STAT3 KO, and IL10 could not inhibit the expression of either miR-155-5p or miR-155-3p in STAT3 KO cells and instead *increased* their levels in STAT3 KO cells. The effects of STAT3 deficiency on LPS and IL10 regulation of miR-155-5p and miR-155-3p can also be seen when the qPCR gene expression data were normalized to each cell type’s LPS-stimulated sample (**Fig 6B**).

**Fig. 6.**
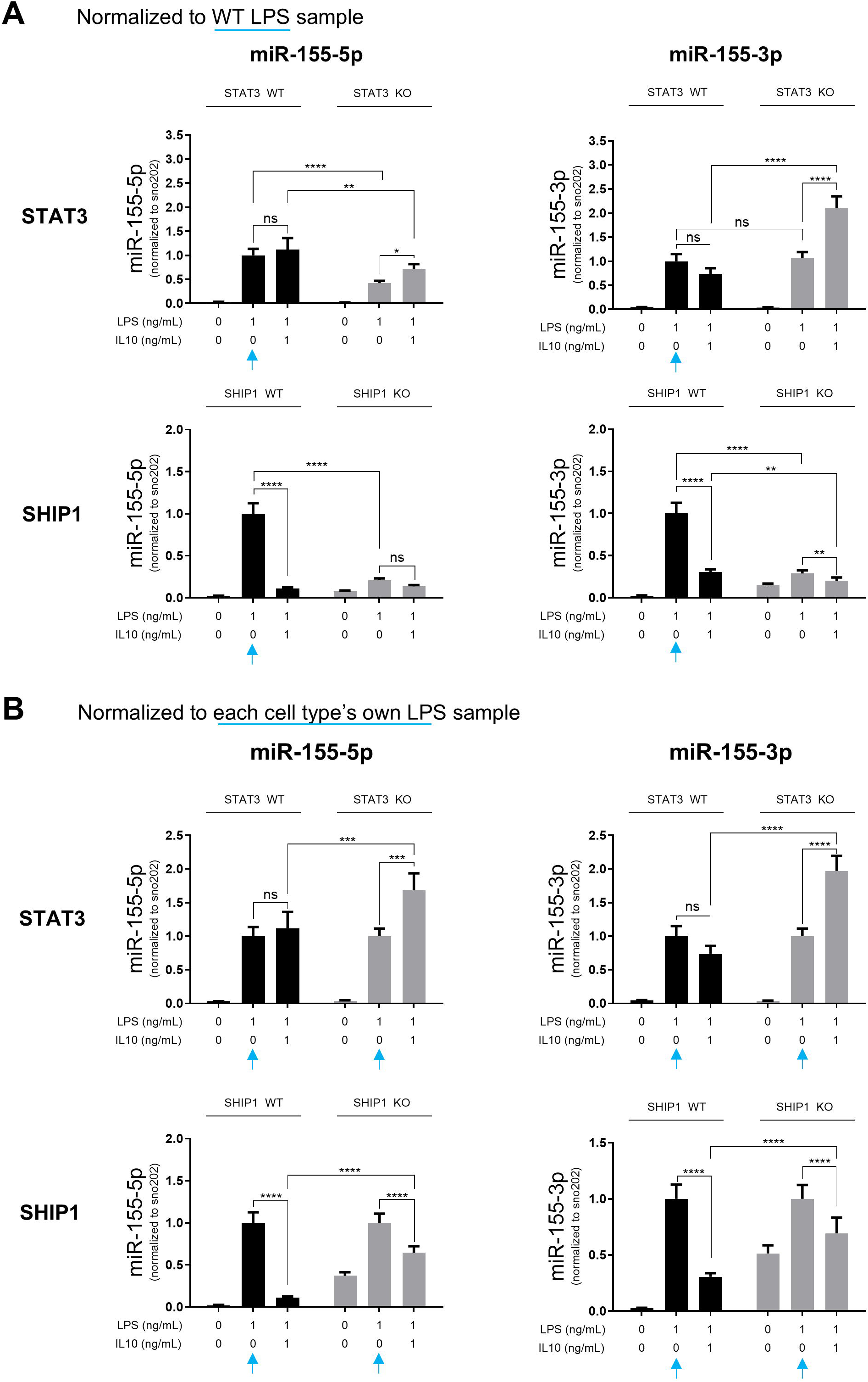
STAT3 and SHIP1 dependence of miR-155-5p and miR-155-3p expression at 4 hours. Expression level in STAT3 WT and KO, or SHIP1 WT and KO cells of miR-155-5p or miR-155-3p was determined by qPCR and normalized to snoRNA202 levels. (A) Data were plotted to represent the RNA expression normalized to the LPS-stimulated sample of the STAT3 WT. (B) The data in panel A was replotted with RNA expression normalized to each cell’s own LPS-stimulated sample. Two-way ANOVA with Tukey’s correction determined the comparison between the stimulations indicated with braces. **** *p* < 0.0001, *** *p* < 0.001, ** *p* < 0.01, ns = not significant. The data are representative of 3 experiments.

For SHIP1, the LPS-stimulated expression of both miR-155-5p and miR-155-3p is decreased in the SHIP1 KO compared to the SHIP1 WT cells (**Fig. 6A**). To see if IL10 could inhibit the LPS-induced expression of miR-155 in the SHIP1 KO, we also normalized the RNA expression data to each cell’s own LPS-stimulated sample. As **Fig. 6B** shows, both miR-155-5p and miR-155-3p levels are inhibited by IL10 in SHIP1 KO cells, but the degree of inhibition is impaired compared to SHIP1 WT. These observations suggest that SHIP1 is required for IL10 inhibition of miR-155-5p and the LPS-induced expression of both miR-155-5p and miR-155-3p.

We then examined the kinetics of miR-155-5p and miR-155-3p expression at 0, 1, 2 and 4 hours (**Fig. 7**). The qPCR data were normalized to the 0 hr time point of the appropriate WT cell control. In STAT3 WT cells stimulated with LPS, miR-155-3p rises rapidly and reaches a plateau at 2 hours, while miR-155-5p rises more slowly but continues to rise towards 4 hours. The presence of IL10 (LPS + IL10) reduced the levels of miR-155-3p but not miR-155-5p in STAT3 WT cells. The different miR-155-5p and miR-155-3p kinetics resemble those observed by Simmonds *et al.* in human peripheral blood-derived macrophages^42^. In STAT3 KO cells, miR-155-5p and miR-155-3p levels stimulated by LPS show similar kinetics to the STAT3 WT. Remarkably, the presence of IL10 (LPS + IL10) *enhanced* rather than inhibited the expression of both miR-155-5p and miR-155-3p. We did not look at longer time points because the effect of autocrine cytokines will contribute to gene expression at longer stimulation times.

**Fig. 7.**
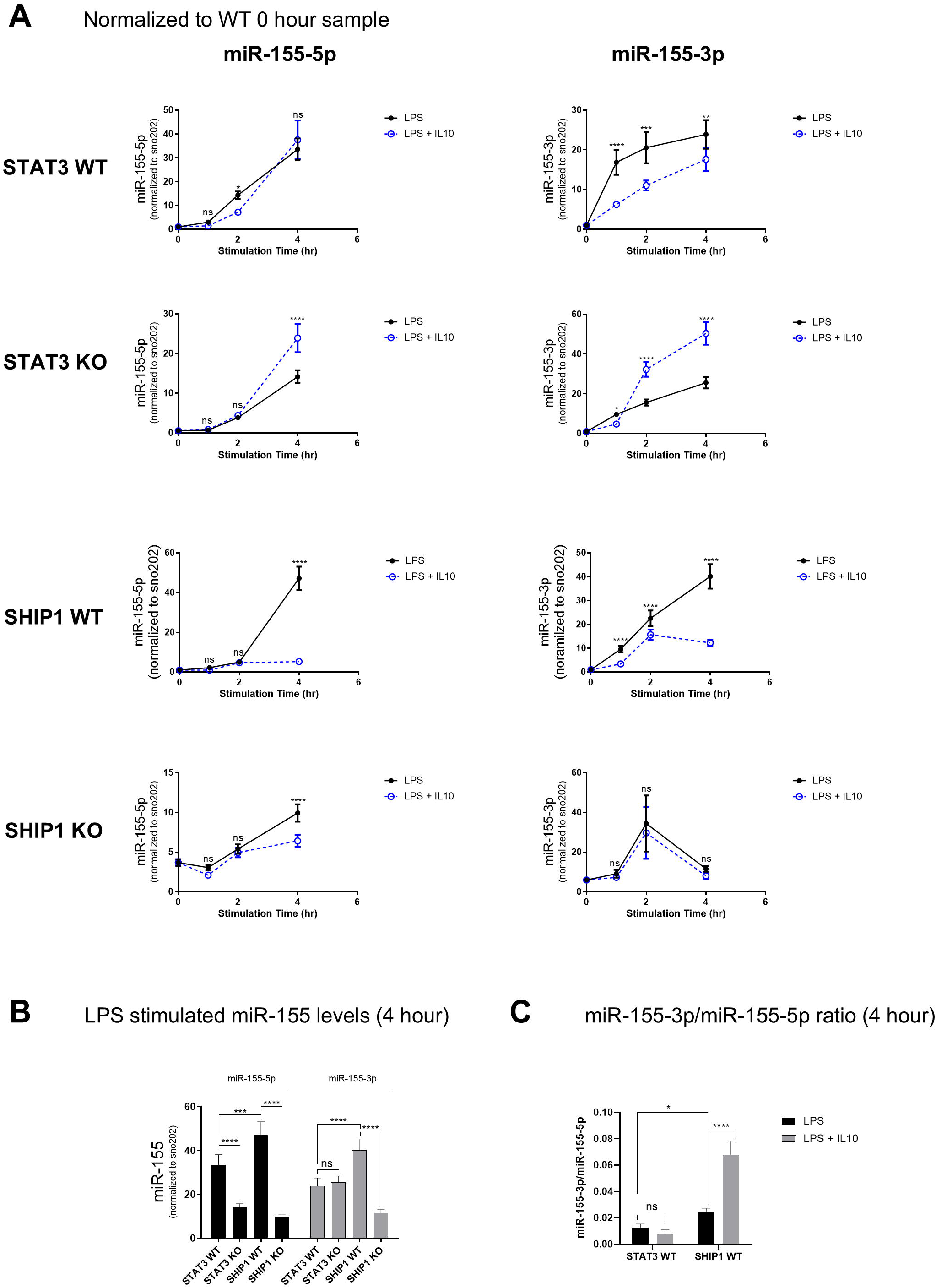
Kinetics of miR-155-5p and miR-155-3p expression in STAT3/SHIP1 WT and KO cells. The expression level in SHIP1 WT and KO cells of miR-155-5p or miR-155-3p was determined by qPCR and normalized to snoRNA202 levels. (A) Kinetics of miR-155-5p and miR-155-3p expression with RNA normalized to the WT 0-hour sample. (B) The data plotted represents the comparison of LPS-stimulated miR-155 levels at 4 hours between different cell types. (C) The data plotted represents the ratio of miR-155-3p and miR-155-5p. Two-way ANOVA with Tukey’s correction determined the significance of the difference in values (if any) in the LPS ± IL10 stimulated sample at the same time point. **** *p* < 0.0001, *** *p* < 0.001, ** *p* < 0.01, * *p* < 0.05, ns = not significant. The data are representative of 3 experiments.

In SHIP1 WT cells, LPS-stimulated miR-155-3p can be detected by 1 hour and continues to rise towards 4 hours. LPS induced high levels of miR-155-5p at 4 hours. In SHIP1 WT cells, IL10 inhibited the LPS-stimulated expression of miR-155-3p and miR-155-5p at all time points. For SHIP1 KO cells, the basal level of both miR-155 strands is elevated, and the LPS-induced miR-155-5p is significantly reduced compared to WT. The LPS-induced miR-155-3p levels are similar to those in SHIP1 WT cells; however, miR-155-3p can no longer be detected at 4 hours. IL10 was impaired in reducing miR-155-5p levels but completely failed to inhibit miR-155-3p levels at all time points.

As **Fig. 7A** shows, the kinetics of both miR-155-5p and miR-155-3p in the SHIP1 WT cells are slightly delayed compared to those in STAT3 WT cells. SHIP1 WT cells also have considerably lower miR-155-5p levels (**Fig. 7B**) compared to STAT3 WT, and an miR-155-3p/miR-155-5p ratio is about twofold (**Fig. 7C**) higher than the STAT3 WT cells. Furthermore, STAT3 WT shows no difference in the ratio of miR-155 strands between LPS and IL10 stimulation, but SHIP1 WT shows a significant increase in the ratio in IL10 stimulation. This difference might reflect the different genetic backgrounds of the SHIP1 WT/KO (Balb/C) and STAT3 WT/KO (C57BL/6) mice.

### The KH3 domains of both FUBP1 and KSRP interact with pre-miR-155

The RNA-binding domains of the FUBP family of RNA-binding proteins have been studied^54^. Previous studies have shown that the RNA-binding protein KSRP interacts with pri-miR-155 and pre-miR-155 to regulate miR-155 expression^25,45^. KSRP and FUBP1 proteins have a conserved architecture of 4 tandem KH domains to interact with RNA/DNA^55^. However, KSRP mainly interacts with RNA via its third KH domain (KH3), and substituting the key GXXG residues of the KH domain with GDDG significantly decreased the interaction of KSRP with RNA^47^. Therefore, to test whether FUBP1 interacts with pre-miR-155 through its KH3 domain, we generated recombinant WT and KH3 domain GDDG mutants for FUBP1 and KSRP (**Fig. 8A**). We then measured the interaction of these recombinant proteins to pre-miR-155 using biolayer interferometry (BLI). **Fig. 8B** and **8C** show that KSRP WT and FUBP1 WT interact with pre-miR-155, but the KH3 GDDG mutants of KSRP and FUBP1 do not.

**Fig. 8.**
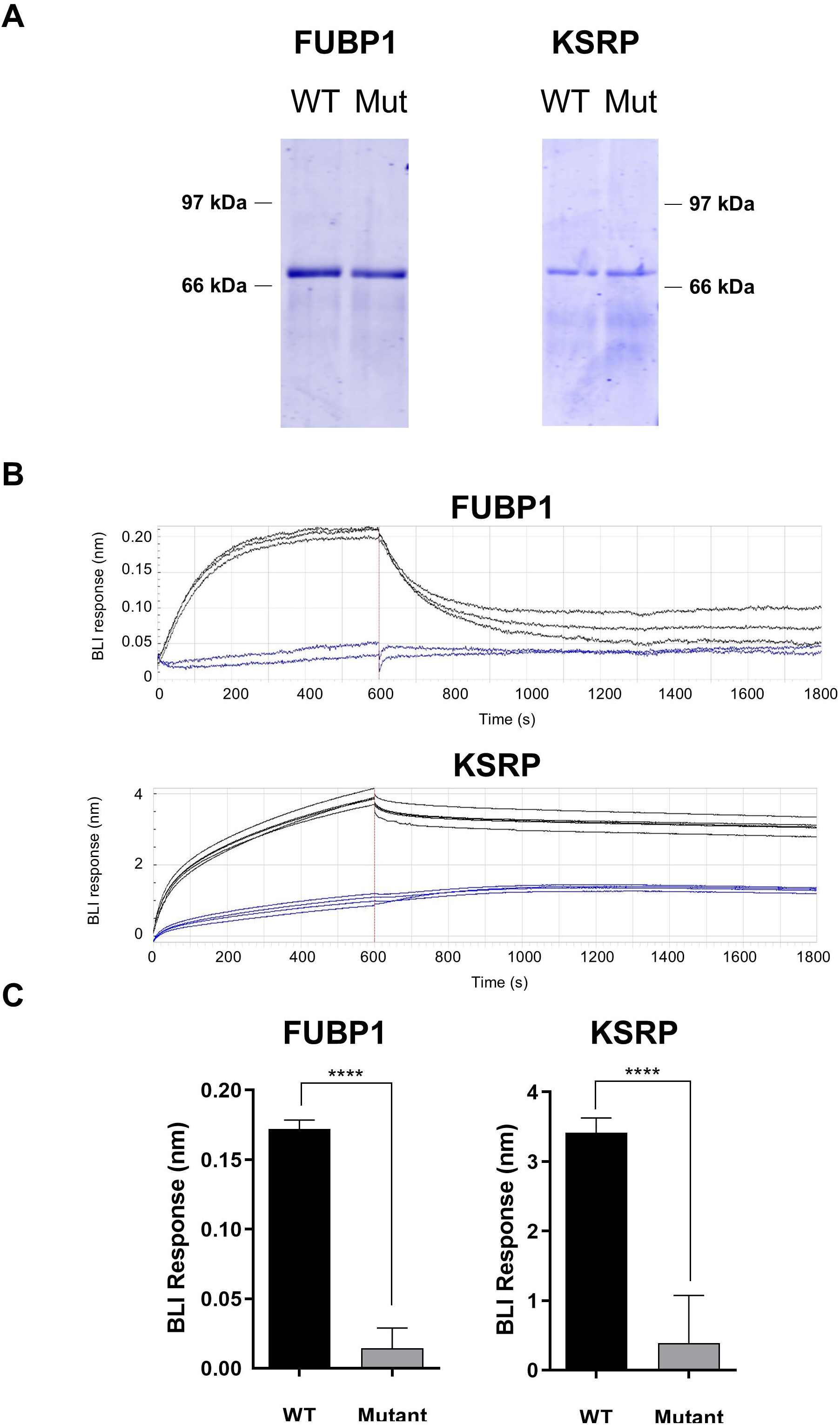
The KH3 domain of KSRP and FUBP1 are essential for interacting with pre-miR-155. KSRP/FUBP1 WT or KSRP/FUBP1 KH3 GDDG mutant proteins were expressed and purified as described in the material and methods. (A) Coomassie Blue staining of the gel assessed the purity of the purified protein. (B) BLI biosensors loaded with biotinylated pre-miR-155 were dipped in wells containing KSRP WT, KSRP KH3 GDDG mutant, FUBP1 WT, FUBP1 KH3 GDDG mutant proteins for 10 minutes, followed by a dissociation step in the assay buffer for 20 minutes. (C) Data plotted to represent the BLI response of indicated proteins with biotinylated pre-miR-155. Unpaired Student’s t-test determined the comparison between the WT and KH3 GDDG mutant. **** *p* < 0.0001. The data are representative of 3-5 experiments.

## Discussion

Most investigators have focused on studying miR-155-5p because it is the dominant miR-155 strand^3,6,7,35^. Simmonds *et al.* were the first to report that LPS-induced miR-155-3p is incorporated in RISC in human macrophages, indicating the non-dominant miR-155-3p is regulated and biologically functional^42^. The miR-155-3p expression peaks at earlier LPS stimulation (4 hours) and disappears by 24 hours^42,45^. In contrast, miR-155-5p levels do not rise until about 2-4 hours and remain detectable in higher numbers at 24 hours^42,45^. Therefore, dissecting the strand-specific control of miR-155-3p and miR-155-5p levels in the cells may provide further information on the mechanisms involved in host immune defence^56^. Moreover, the dysregulation of miR-155-3p has been reported in various cancer and disease cells. The dysregulated level of miR-155-3p in the cancer and disease cell may be due to increased expression of the *MIR155HG* gene^57^, increasing both miR-155-5p and miR-155-3p strands^58–61^. However, there are also cases where only the miR-155-3p is upregulated in the cancer and disease cells, indicating strand-specific control of miR-155-3p^13,15,62,63^. Furthermore, the two strands of miR-155 have different targets but overlapping targets^58,59^. Therefore, the ratio of the strands expressed and target availability may be important factors determining the overall effects of miR-155 dysregulation on cancer growth^64^ and diseases^65^.

### Role of CELF2, FUBP1 and KSRP in controlling expression of miR-155-5p and miR-155-3p

Our examination of the CELF2, FUBP1, and KSRP KD cells revealed different miR-155-5p and miR-155-3p expression patterns compared to the control RAW264.7 cells. **(1)** Without CELF2 protein, LPS-induced expression of miR-155-5p increases significantly compared to control RAW264.7 cells at 2 and 4 hours. In contrast, miR-155-3p is modestly increased at 2 hours and decreased compared to control RAW264.7 cells at 4 hours. The resulting decrease in the miR-155-3p/miR-155-5p ratio suggests that the CELF2 protein plays a role in strand switching to miR-155-3p from miR-155-5p (**Fig. 3**). **(2)** Knocking down FUBP1 protein increased miR-155-3p expression, but the overall cellular ratio of miR-155-3p/miR-155-5p remains unchanged. This observation suggests that the FUBP1-mediated miR-155-3p molecules may have subcellular compartment/location-specific functions, as reported for other miRNAs^66^. **(3)** The absence of KSRP protein resulted in a substantial increase in the levels of both strands of miR-155. KSRP may thus be a general inhibitor of the miR-155 maturation process.

Our previous studies showed that the CELF2 protein inhibits miR-155 maturation^37^, similar to the CELF2 function on miR-140 maturation^41^. However, our current studies showed that the CELF2 protein differentially regulates the miR-155 strands (**Fig. 3** and **4**). Without CELF2 protein, miR-155-5p is enhanced, but miR-155-3p is reduced (**Fig. 3** and **4**). We have also looked at the levels of the primary transcript of miR-155 (pri-miR-155) and the precursor of miR-155 (pre-miR-155) in these cells (**Supplementary Fig. 1**). The pri-miR-155 and pre-miR-155 levels in CELF2 KD cells were indifferent to the control cell, stating that the change in miR-155 levels occurs at the stage of maturation of miR-155. These observations suggest that the CELF2 protein regulates miR-155-3p expression and is an essential strand-switching factor, choosing miR-155-3p over miR-155-5p at early time points by LPS stimulation.

CELF2 protein has been described to possess anti-tumour functions, where low expression of CELF2 protein was detected in various cancer cells. Overexpression of CELF2 showed a reduction in the size of the tumour cells^67–70^. Interestingly, the levels of miR-155-5p and miR-155-3p in cancer cells have been reported to be predominantly upregulated^58,60^. A higher level of CELF2 protein may explain the high level of miR-155-3p in these cancer cells ^53,58,60,64^. Therefore, future studies investigating the correlation between the miR-155-5p and miR-155-3p levels and CELF2 expression levels in various cancer types will provide more information on miR-155 regulation and how CELF2 protein contributes to it.

Ruggiero *et al.* reported that KSRP binds pre-miR-155 and is required to process pre-miR-155 to miR-155-5p^25^. They showed that in the absence of KSRP protein, miR-155-5p levels are reduced, and miR-155-3p levels are increased, proposing the role of KSRP as a selective miR-155-5p regulator^25^. Our work partly agrees with data from Ruggiero *et al.,*as KSRP KD shows an increased miR-155-3p level. However, our data show increased miR-155-5p levels at 4 hours of LPS stimulation. Our work suggests that the KSRP protein functions as a general inhibitor of the maturation of miR-155. Thus, both strands of miR-155 are increased in KSRP KD cells. The discrepancy in data may be due to the difference in concentration of LPS used or the different methods of generating the knockdown cell lines. We used a physiologically relevant concentration of 1 ng/mL LPS, while Ruggiero *et al.* used the super-physiological concentration of 100 ng/mL of LPS. Very high ligand concentrations can change the kinetics of gene expression and/or set up feedback effects not seen at lower ligand concentrations. Further work is underway to examine these possibilities.

FUBP1 has not been previously described to participate in miRNA processing. Based on our work, the role of FUBP1 in miR-155 regulation is specific to miR-155-3p (**Fig. 3**). Furthermore, our work indicates the KH3 domain of FUBP1 participates in binding to pre-miR-155, analogous to the requirement of the KSRP KH3 domain for miR-155-5p binding^25^. The KH3 domain of FUBP1 and KSRP share 79% sequence homology (**Fig. 11A**). Thus, KSRP and FUBP1 may recognize the same or similar sequences in pre-miR-155. The consensus recognition motif for the KSRP KH3 domain (GGGG)^71^ and FUBP1 KH3 domain (UUGUG)^72^ both appear at the 3′ end of the miR-155-5p sequence of pre-miR-155 (**Fig. 11B**). Interestingly, the sequence recognition motif for CELF2 protein (UGU-N_x_-UGU)^73^ exactly overlaps two of three expected FUBP1 recognition motifs in the pre-miR-155 sequence, suggesting the possibility of FUBP1 to compete for CELF2 instead of KSRP protein.

### IL10R signaling control of miR-155-5p and miR-155-3p expression

As a next step in investigating how these three proteins coordinate their control of miR-155-5p and miR-155-3p, we also examined the IL10-regulated SHIP1 and STAT3 dependence of miR-155-5p and miR155-3p expression IL10 signalling downstream of the IL10R involves both STAT3 and SHIP1:STAT3 pathways^31^. Our studies showed that in the absence of STAT3 protein, IL10 no longer inhibits either miR-155-5p or miR-155-3p production but instead enhances expression of both by 4 hours and 2 hours, respectively (**Fig. 7A**). In the absence of SHIP1, the LPS-induced level of both miR-155 strands is significantly reduced, IL10-dependent inhibition of miR-155-5p is partly impaired and IL10-dependent inhibition of miR-155-3p is abolished (**Fig. 6B**). Our data suggests that STAT3 and SHIP1 underlie miR-155 regulation by the IL10R pathway, as summarized in **Fig. 9**. We found in WT mouse macrophages that LPS-induced miR-155-3p expression increases more quickly than miR-155-5p (**Fig. 7**). In the C57BL/6 mouse (STAT3 WT), miR-155-3p reaches a plateau at around 2 hours. In the Balb/C mouse (SHIP1 WT), the miR-155-3p level continuously rises to 4 hours. The difference in miR-155-3p kinetics between STAT3 WT and SHIP1 WT may be due to the difference in mouse strain (C57BL/6 vs Balb/C) and may contribute to the described differing immune response in these strains ^74,75^.

**Fig. 9.**
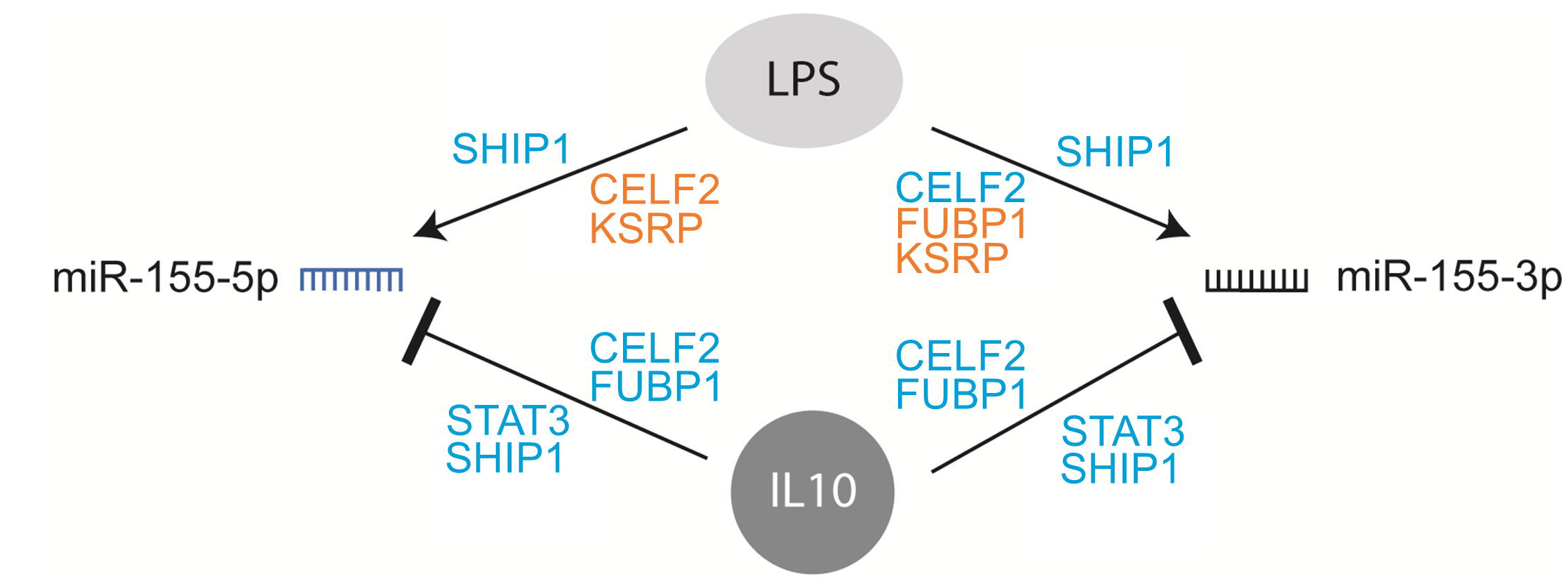
Schematic representation of miR-155 regulation. Schematic diagram representing the regulation of miR-155-5p and miR-155-3p expression in response to LPS and IL10. The proteins in **blue** font represent the protein **required** for the shown action, and the proteins in **orange** font represent proteins that **suppress** the shown action.

We had previously examined the kinetics of IL10 inhibition of LPS-induced expression of TNFα^31^. We showed that the early (<2 hours) response required a SHIP1:STAT3 complex, while the late phase (>2 hours) only needed STAT3. In the current study, we support the previous findings^42^ of miR-155-3p kinetics by LPS being earlier (2-4 hours) than miR-155-5p (>4 hours). Furthermore, in the absence of STAT3 or SHIP1 protein, IL10-dependent inhibition of miR-155-3p is abolished, whereas, for miR-155-5p, SHIP1 KO only resulted in partial impairment of IL10-dependent inhibition of miR-155-3p (**Fig. 7A**). These observations suggest that IL10 inhibition of miR-155-3p may be necessary for IL10 inhibiting the early (SHIP1:STAT3 dependent) phase of TNFα production. In contrast, miR-155-5p inhibits the late (STAT3-dependent) phase of TNFα expression. However, the mechanism by which STAT3 or SHIP1:STAT3 complexes control miR-155-5p and miR-155-3p remains to be determined. One possibility is that both lead to protein expression that participates in the control of pre-miR-155 processing. In fact, since the absence of STAT3 results in IL10 *enhancing* rather than *reducing* LPS-induced miR-155-5p and miR-155-3p levels, we predict these STAT3-induced proteins may suppress pre-miR-155 processing (**Fig. 10**).

**Fig. 10.**
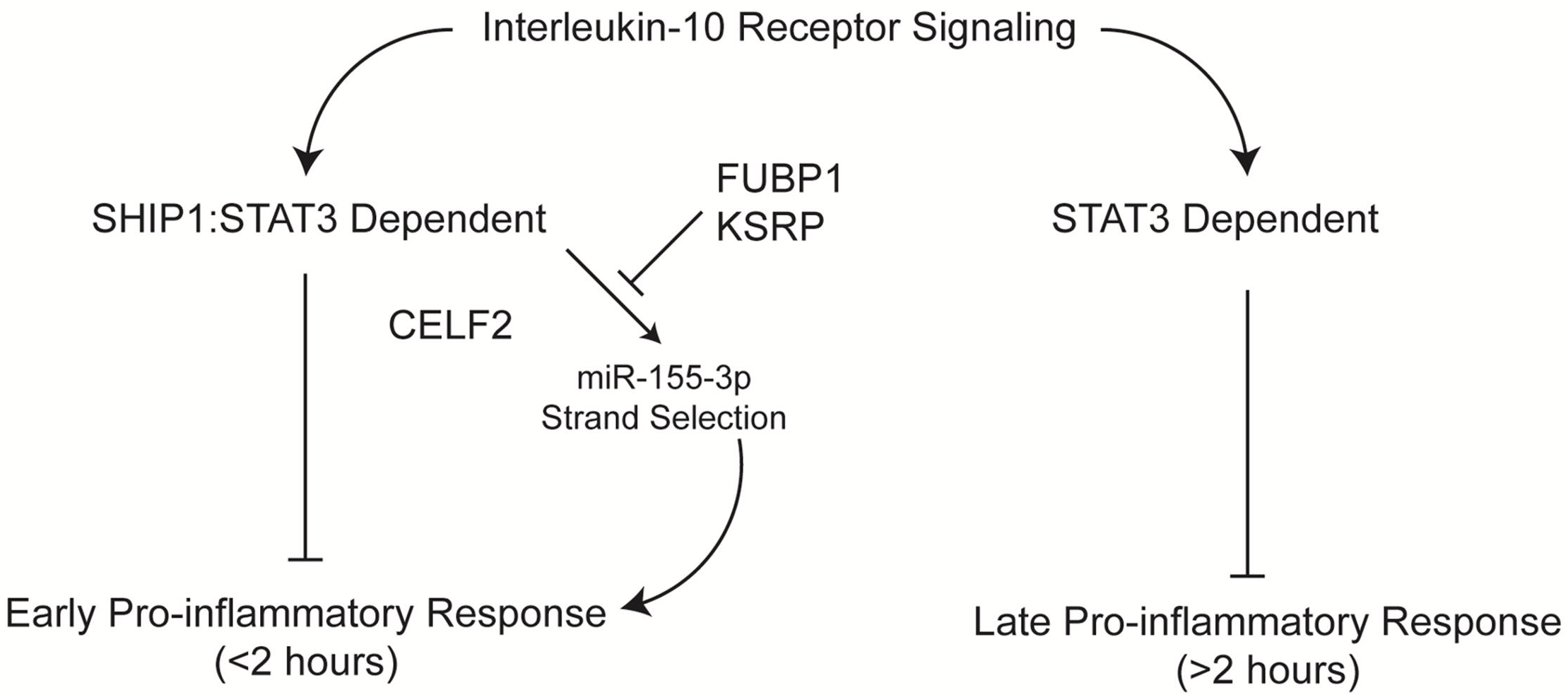
Schematic representation of IL10R downstream signaling process and potential mechanism on a different phases of pro-inflammatory response. Schematic diagram of IL10R downstream signaling pathway affecting various stages of pro-inflammatory response.

**Fig. 11.**
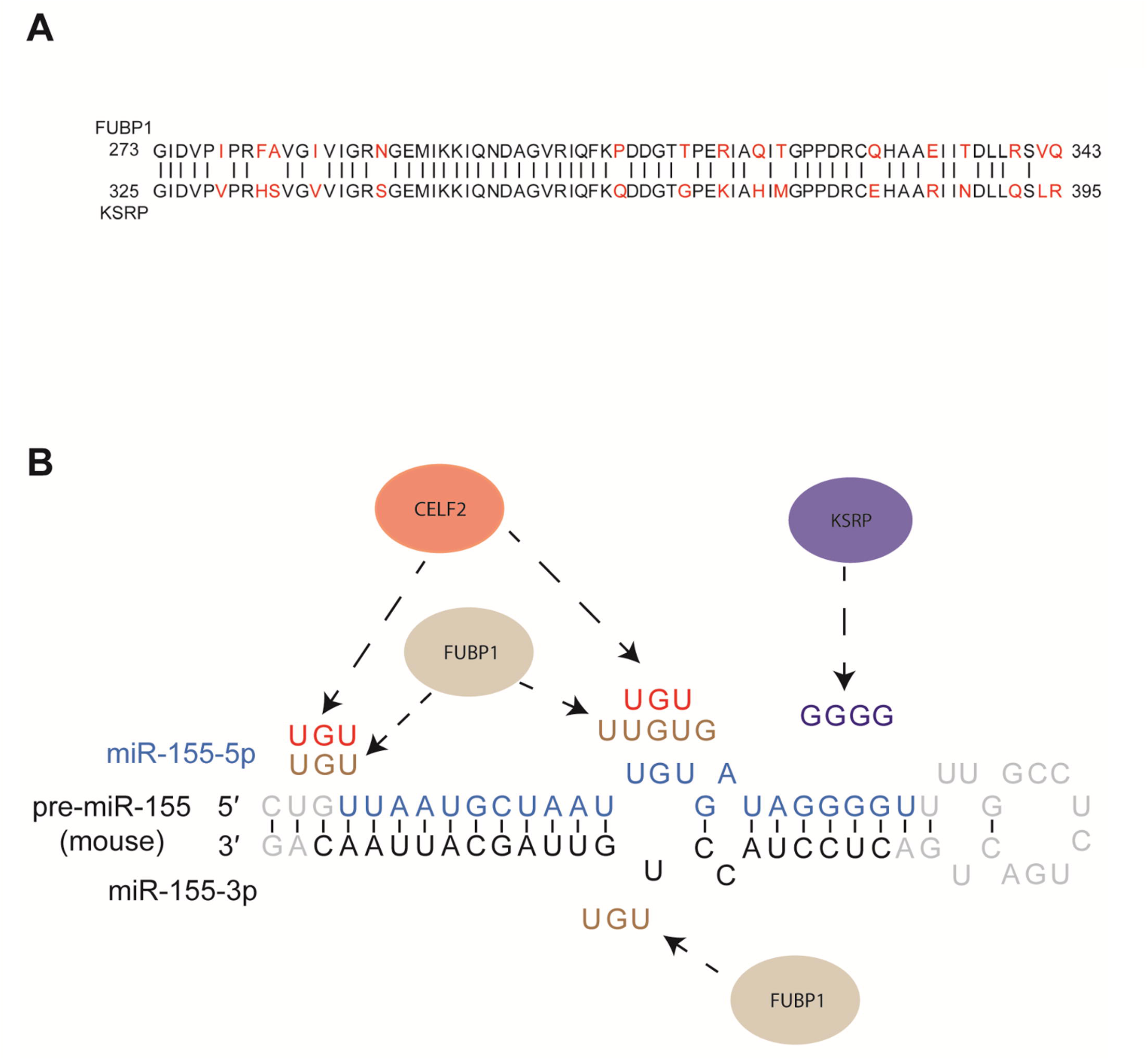
Schematic representation of FUBP1/KSRP /CELF2 regulation, homology, and interaction. (A) The sequence alignment of the KH3 domain of FUBP1 and KSRP protein. Sequences in black and red indicate matching and mismatching sequences, respectively. (B) Schematic diagram showing the predicted interaction site of FUBP1, CELF2 and KSRP to pre-miR-155. The bases of pre-miR-155 are in gray; blue bases show the miR-155-5p sequence, and black bases indicate the miR-155-3p sequence.

Work is underway to determine how CELF2, SHIP1, and STAT3-dependent pathways affect miR-155 strand selection. Two strands of miRNA can be produced from a single precursor molecule of miRNA, but only one strand is selected to load into the RNA-induced silencing complex^76^. Strand selection of miRNAs may occur due to a change in the cleavage site of Dicer^77^ or through actions of RNA editing enzymes on the pri/pre-miR-155 molecule^76^ that affects the thermodynamic stability of the strands or its interaction with RNA binding proteins. Altered thermodynamic stability or Dicer cut site may result in selective processing of one or the other strand into the one that becomes the guide strand^78^. In the case of miR-155, the CELF2 protein shows the characteristics of strand selection, favouring the miR-155-3p expression and FUBP1 favouring miR-155-5p expression. We will also investigate whether RNA editing participates in miR-155-5p vs miR-155-3p expression and whether these participate in the aberrant expression of miR-155-5p vs miR-155-3p observed in certain cancers^53,58,60,64^. These investigations will also provide insight into regulating other miRNAs’ expression, including those described as regulated by CELF2 and KSRP^41,44,79–81^.

## Materials and Methods

### Mouse colonies

SHIP1 WT or SHIP1 KO (in the BALB/c background) mice were provided by Dr. Gerald Krystal (BC Cancer Research Centre, Vancouver, BC). STAT3 WT and KO mice (in the C57BL/6 background) were generated as described^31^. All mice were maintained following the animal care protocols approved by the University of British Columbia Animal Care Committee.

### Cell lines

The murine cell line RAW264.7 (ATCC TIB-71) was maintained in Roswell Park Memorial Institute 1640 medium (RPMI-1640, SH30027, HyClone, Logan, UT) supplemented with 9% fetal bovine serum (FBS, SH30396, HyClone, Logan, UT). As described below, FUBP1 KD RAW264.7 cells were generated by transduction of RAW 264.7 expressing iCas9 (called RAW264.7 control cells). The generation of CELF2 KD RAW264.7 cells was described in Yoon *et al.*^37^.

### Construction of the FUBP1 or KSRP-pLX-sgRNA targeting vectors

The FUBP1 sgRNA targeting sequence (5’ GCTAAATCCGACCATCCCATC) was designed using the CRISPR Gold online tool^82^. Oligonucleotides corresponding to this sequence were cloned into the pLX-sgRNA vector using overlap-extension PCR as described^37^. The pLX-sgRNA vector with target-specific insert was transformed into chemically competent Stbl3 *E. coli* cells, and colonies were selected using ampicillin. The resulting FUBP1-pLX-sgRNA construct was confirmed by sequencing.

### Generation of RAW 264.7 cells expressing FUBP1 or KSRP pLX-sgRNA

The FUBP1-pLX-sgRNA vector was transduced into RAW264.7 cells expressing iCas9 (called RAW264.7 control cells)^37^ using lentiviruses. Lentiviruses harbouring FUBP1-pLX-sgRNA were prepared by co-transfecting FUBP1-pLX-sgRNA vector into HEK293T (ATCC CRL-3216) cells with the packaging plasmid R8.9 and VSVG. 24 hours after transfection, the supernatant was collected and incubated with RAW264.7 control cells in the presence of 8 µg/mL protamine sulfate. Cells transduced with FUBP1-pLX-sgRNA viruses were selected using 10 µg/mL blasticidin2 µg/mL doxycycline was added to the culture media for up to 48 hours to induce the expression of Cas9 and knockdown of FUBP1.

### RNA-oligonucleotide transfection and RNA pull-down assay

RAW264.7 cells (ATCC TIB-71) were seeded at 8.4 × 10^6^ cells per 10 cm dish a day before the transfection. Biotinylated pre-miR-155 oligonucleotides (Biotin-UAAUUGUGAUAGGGGUUU UGGCCUCUGACUGACUCCUACCUGUUA) were obtained from Invitrogen Life Technologies (ThermoFisher Scientific, Nepean, ON). RNA oligonucleotides and Lipofectamine-3000 (L3000-015, ThermoFisher Scientific, Nepean, ON) were prepared in Opti-MEM (31985, ThermoFisher Scientific, Nepean, ON) separately. They was mixed at 1:1 (v/v) ratio, incubated at room temperature for 20 minutes to allow the formation of Lipofectamine-oligonucleotide complexes. 1 µg of RNA-oligonucleotide to 1.125 µL of Lipofectamine-3000 was used. After incubation, the RNA-Lipofectamine solution was diluted ten-fold with 9% FBS/RPMI-1640 and added to cells. The cells were incubated with the transfection solution in a chamber at 37°C supplemented with 5% CO_2_. After 6 hours, the solution was replaced with 9% FBS/RPMI-1640, and cells were allowed to recover overnight

Following the 9% FBS/RPMI-1640 overnight incubation, the transfected RAW264.7 cells were stimulated with 1 ng/mL LPS (*Escherichia coli* serotype 0111:B4; Millipore-Sigma, Oakville, ON) ± 50 ng/mL IL10 for the indicated length of time in a chamber at 37°C supplemented with 5% CO_2_. Following stimulation, the media was removed, and cells were chilled with cold (4°C) phosphate-buffered saline (PBS, SH30256, ThermoFisher Scientific, Nepean, ON) for 2 minutes. Next, the PBS was removed, and the cells lysed by the addition of protein solubilization buffer (PSB, 50 mM HEPES, 100 mM NaF, 10 mM NaPPi, 2 mM NaVO_4_, 2 mM NaMoO_4_, 4 mM EDTA, 0.125% Triton X-100, protease inhibitor cocktail (11836145001, Millipore-Sigma, Oakville, ON), and 0.5 mM Tris(2-carboxyethyl)phosphine (TCEP, M115, Soltec Ventures, Beverly, MA)). Cell lysates were collected with a cell scraper and gently agitated at 4°C for 30 minutes. Insoluble material was removed by centrifugation at 12,000 RPM for 20 minutes at 4°C.

Clarified cell lysates were added to streptavidin magnetic beads (1164786001, Millipore-Sigma, Oakville, ON) in 1.5 mL microfuge tubes and incubated for 90 minutes at 4°C on a nutator. The tubes were then briefly centrifuged at 5,000 RPM, and magnetic beads were immobilized using a magnetic tube stand (12321D, ThermoFisher Scientific, Nepean, ON). Lysates were removed, and the beads were resuspended in the wash buffer (0.1% Tween-20 containing PSB) and rocked for 5 minutes at 4°C on a nutator. The washing was repeated 3 times. The proteins were eluted by boiling 2x SDS-PAGE sample buffer (0.125 M Tris, pH 6.8, 5% 2-mercaptoethanol, bromophenol blue, 13.5% glycerol, 4.5 % SDS) for immunoblot analysis.

### Immunoblot analysis

Proteins were separated by 10% SDS-PAGE, followed by electroblotting onto polyvinylidene fluoride (PVDF) membrane (IPFL00010, Millipore-Sigma, Oakville, ON). The membranes were blocked in 3% bovine serum albumin (BSA), then probed with the following primary antibodies overnight: 1:1000 KSRP (ab140648, Abcam, Toronto, ON), 0.1 µg/mL GAPDH (G9545, Millipore-Sigma, Oakville, ON), 1 µg/mL Myd88(Sc-74532, Santa Cruz, Dallas, TX), 1 µg/mL Actin (A2066, Sigma, Oakville, ON) and 1 µg/mL FUBP1 or KSRP (Sc-136137, Santa Cruz, Dallas, TX). The membranes were washed three times in Tris-buffered saline containing 0.05% Tween-20 (TBST), incubated with either Alexa Fluor 660 anti-mouse IgG (A21055) or Alexa Fluor 680 anti-rabbit IgG (A21109, Invitrogen, Burlington, ON), and imaged using a LI-COR Odyssey Imager.

### Isolation of and stimulation of mouse peritoneal macrophages

Primary peritoneal macrophages (perimacs) were isolated from mice by peritoneal lavage with 3 mL of sterile PBS. Perimacs were seeded at 2.0 × 10^6^ cells per well in a 6-well tissue culture plate or 1.18 × 10^6^ cells per well in a 24-well tissue culture plate in Iscove’s Modified Dulbecco’s Medium (IMDM, SH30228, HyClone, Logan, UT) supplemented with 10% FCS. Cells were allowed to adhere for 2 hours, rinsed with room temperature PBS to remove non-adhered cells, and fresh media was added. The media was changed after 1 hour, and the cells were stimulated with 1 ng/mL LPS ± 1 ng/mL IL10 for 1, 2, or 4 hours. Triplicate wells were used for each stimulation condition.

### RNA extraction and qPCR

Total RNA was extracted using Tri-Reagent (T9424, Millipore-Sigma, Oakville, ON) according to the manufacturer’s instructions. 1-3 µg of RNA was treated with RNase-free DNase I (04716728001, Millipore-Sigma, Oakville, ON) for 20 minutes at 37°C, followed by the addition of 0.1 M EDTA to a final concentration of 8 mM to inactivate DNase I.

For measurement of miR-155-5p, miR-155-3p, and small nucleolar RNA MBII-202 (snoRNA202) levels, 20 ng of DNase I treated RNA was used to generate cDNAs using the miRNA TaqMan Reverse Transcription Kit (4366597, ThermoFisher Scientific, Nepean, ON), Multiscribe™ reverse transcriptase (4319983, ThermoFisher Scientific, Nepean, ON), and miR-155-5p (002571), miR-155-3p (464539_mat) or snoRNA202 (001232, ThermoFisher Scientific, Nepean, ON) primers according to the manufacturer’s instructions. qPCR quantification of miR-155-5p and miR-155-3p and snoRNA202 cDNA was performed using the TaqMan fast advanced master mix (4444557, ThermoFisher Scientific, Nepean, ON) and the appropriate TaqMan probes on a StepOne Plus™ instrument (4376582, Invitrogen, Burlington, ON). miRNA levels were analyzed using the comparative CT method with snoRNA202 as the normalization control.

For measurement of pri-miR-155, pre-miR-155, and GAPDH, 200 ng of DNase I treated RNA were reverse transcribed using SuperScript™ IV reverse transcriptase (18090200, ThermoFisher Scientific, Nepean, ON), oligodT primer (18418020, ThermoFisher Scientific, Nepean, ON) and random hexamers. qPCR quantification of pri-miR-155, pre-miR-155, and GAPDH were achieved with primers for pri-miR-155, pre-miR-155, and GAPDH in conjunction with the SYBR Green master mix (100029284, ThermoFisher Scientific, Nepean, ON). RNA levels were analyzed using the comparative CT method with GAPDH as the normalization control.

### Molecular cloning of FUBP1 and KSRP

The open reading frame (ORF) of mouse FUBP1 and KSRP was obtained by PCR on cDNA generated from RNA isolated from SHIP1 WT perimacs. The FUBP1/KSRP ORF was inserted into the Gateway entry vector (pENTR1A, Invitrogen, Burlington, ON) via restriction digest and ligation. The products were transformed into DH5α *E. coli* chemically competent cells, and colonies were selected using 50 µg/mL of kanamycin. The sequences of the FUBP1/KSRP pENTR1A vectors were confirmed by sequencing. Using site-directed mutagenesis, the GXXG sequences in the third KH domain of FUBP1 and KSRP were mutated to the GDDG sequence. PCR with non-overlapping primers (5′ phosphorylated forward primer with desired mutation and non-overlapping reverse primer) was used to generate FUBP1/KSRP daughter plasmids containing the desired mutation. The resulting PCR product was extracted with phenol-chloroform, treated with Dpn I to remove the parental vector, daughter plasmids ligated with T4 ligase, and transformed into DH5α chemically competent cells. The FUBP1/KSRP WT and KH3 GDDG mutant pENTR1A vectors were transferred to lentiviral vector FUGWBW using Gateway LR reactions as described^31^.

### Recombinant FUBP1 and KSRP Protein Expression

HEK293T cells were transfected with FUBWBW vectors encoding FUBP1/KSRP WT or KH3 GDDG mutant proteins. After transfection (48 hours), the cells were lysed with the lysis buffer (20 mM Tris-HCl, pH 7.4, 150 mM NaCl, 5 mM imidazole, 1% NP-40, protease inhibitor cocktail, and 10 mM TCEP). The lysates were incubated for 45 minutes at 4°C on a nutator, then centrifuged at 12000 rpm for 20 minutes, and the supernatants were transferred to tubes containing cobalt affinity beads (TALON metal affinity resin, #635504, Takara Bio, Mountain View, CA). The cell lysates were incubated with the beads for 2 hours, and the beads were washed three times with wash buffer (20 mM Tris-HCl, pH 7.4, 150 mM NaCl, 5 mM Imidazole, 0.1% NP-40, 0.5 mM TCEP) before eluting with the elution buffer (20 mM Tris-HCl, pH 7.4, 150 mM NaCl, 150 mM imidazole, 0.5 mM TCEP, and 0.02% Tween-20).

### Measurement of TNF**α** production

Cells were seeded at 2.0 × 10^4^ cells per well in a 96-well tissue culture plate and allowed to adhere overnight. The media was changed the next day, 1 hour before stimulation. Cells were stimulated with 1 ng/mL of LPS ± indicated concentration of IL10 for 1 hour. Triplicate wells were used for each stimulation condition. The supernatant was collected, and secreted TNFα protein levels were measured using a BD OptEIA Mouse TNFα Enzyme-Linked Immunosorbent Assay (ELISA) kit (558534, BD Bio-sciences, Mississauga, ON).

### Biolayer interferometry

The binding affinity between the FUBP1 or KSRP proteins to pre-miR-155 was examined using biolayer interferometry with super-streptavidin (SSA) biosensor tips (18-5057, ForteBio, Fremont, CA). SSA biosensor tips were hydrated in BLI assay buffer (20 mM Tris-HCl, pH 7.4, 150 mM NaCl, 0.5 mM TCEP, 0.2% Tween-20) before coating with biotinylated pre-miR-155 and blocking with 0.1% BSA. The kinetic measurements were done at 30°C with an orbital flow of 1000 rpm. A 60-second baseline was established using BLI assay buffer. The pre-miR-155 coated biosensors were then dipped into wells containing KSRP or FUBP1 protein, and association was monitored for 600 seconds. The sensors were then transferred to wells containing only BLI assay buffer, and protein dissociation was monitored for 600 seconds. The raw data were analyzed using the Octet Red Data Analysis software (ver. 8.2).

### Statistical analysis

Band intensities were quantified in immunoblots using LI-COR Odyssey imaging system and Image Studio™ Lite software (LI-COR Biosciences, Lincoln, NE). Graph-Pad Prism 6 (GraphPad Software Inc., La Jolla, CA) performed all statistical analyses. Statistical details can be found in figure legends. Values are presented as means ± standard deviations. Unpaired Student’s *t*-tests were used where appropriate to generate two-tailed P values. Two-way ANOVA was performed where required with appropriate multiple comparison tests. The differences were considered significant when *p* ≤ 0.05.

### Ethical statement

The perimacs were derived from mice in accordance with the animal care protocol (A21-0203) approved by the University of British Columbia Animal Care Committee. The cell culture experiments are done in accordance with UBC Biosafety requirements (B16-0206).

## Data availability statement

All data not provided in the manuscript and the supplementary information are available from the authors upon request.

## Acknowledgements

We thank Dr. Nada Lallous from Vancouver Prostate Centre for providing help with the BLI experiments and Dr. Gerald Krystal for the SHIP1 WT and KO mice. We also thank Mike K. Wu for their advice on the RNA pull-down experiments.

## Author contributions

Conceptualization, J.Y., and A.M.; methodology, J.Y.; validation, J.Y., T.C. and A.M.; formal analysis, J.Y.; investigation, J.Y., A.W., and A.M.; Resources, J.Y., and T.C.; data curation, J.Y., and A.W..; writing—original draft preparation, J.Y., and A.W.; writing— review and editing, J.Y., T.C. and A.M.; visualization, J.Y.; supervision, A.M.; project administration, T.C., and A.M.; funding acquisition, A.M. All authors have read and agreed to the published version of the manuscript.

