## Supplementary Figure 1 for "The strand-selective regulation of miR-155 in response to lipopolysaccharide by CELF2, FUBP1 and KSRP proteins"

### Slide 1
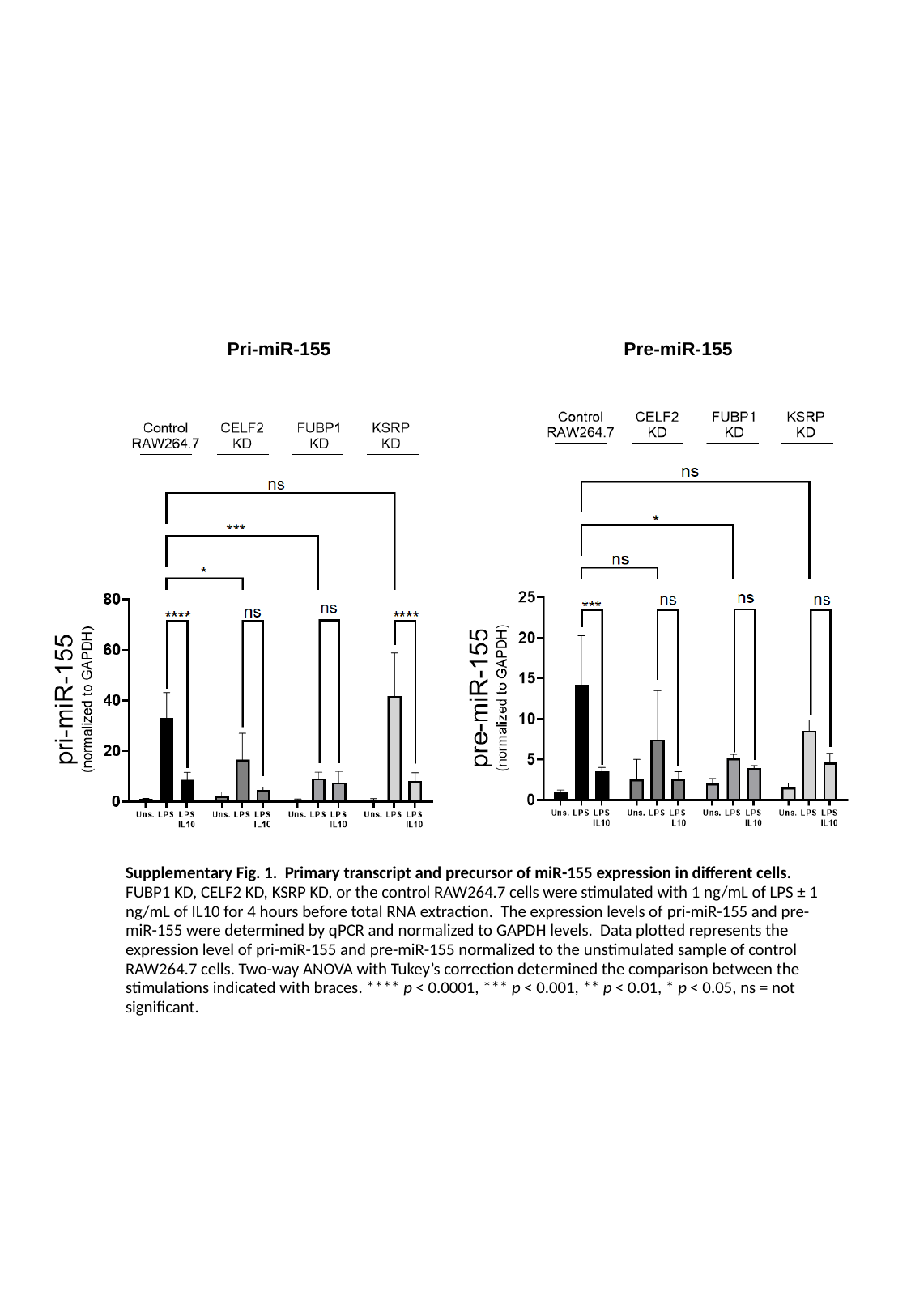

Pri-miR-155
Pre-miR-155
Supplementary Fig. 1. Primary transcript and precursor of miR-155 expression in different cells. FUBP1 KD, CELF2 KD, KSRP KD, or the control RAW264.7 cells were stimulated with 1 ng/mL of LPS ± 1 ng/mL of IL10 for 4 hours before total RNA extraction. The expression levels of pri-miR-155 and pre-miR-155 were determined by qPCR and normalized to GAPDH levels. Data plotted represents the expression level of pri-miR-155 and pre-miR-155 normalized to the unstimulated sample of control RAW264.7 cells. Two-way ANOVA with Tukey’s correction determined the comparison between the stimulations indicated with braces. **** p < 0.0001, *** p < 0.001, ** p < 0.01, * p < 0.05, ns = not significant.
